# Label-free Raman microspectroscopy for identifying virocells

**DOI:** 10.1101/2021.09.22.461451

**Authors:** Indra Monsees, Victoria Turzynski, Sarah P. Esser, André Soares, Lara I. Timmermann, Katrin Weidenbach, Jarno Banas, Michael Kloster, Bánk Beszteri, Ruth A. Schmitz, Alexander J. Probst

## Abstract

Raman microspectroscopy has been thoroughly used to assess growth dynamics and heterogeneity of prokaryotic cells. Yet, little is known about how the chemistry of individual cells changes during infection with lytic viruses, resulting in so-called virocells. Here, we investigate biochemical changes of bacterial and archaeal cells of three different species in laboratory cultures before and after addition of their respective viruses using single-cell Raman microspectroscopy. By applying multivariate statistics, we identified significant differences in the spectra of single cells and cells after addition of lytic phage (*phi6*) for *Pseudomonas syringae*. A general ratio of wavenumbers that contributed the greatest differences in the recorded spectra was defined as an indicator for virocells. Based on reference spectra, this difference is likely attributable to an increase in nucleic acid vs. protein ratio of virocells. This method proved also successful for identification of *Bacillus subtilis* cells infected with *phi29* displaying a decrease in respective ratio but failed for archaeal virocells (*Methanosarcina mazei* with Methanosarcina Spherical Virus) due to autofluorescence. Multivariate and univariate analyses suggest that Raman spectral data of infected cells can also be used to explore the complex biology behind viral infections of bacteria. Using this method, we confirmed the previously described two-stage infection of *P. syringae*’s *phi6* and that infection of *B. subtilis* by *phi29* results in a stress response within single cells. We conclude that Raman microspectroscopy is a promising tool for chemical identification of Gram-positive and Gram-negative virocells undergoing infection with lytic DNA or RNA viruses.

**Importance:** Viruses are highly diverse biological entities shaping many ecosystems across Earth. Yet, understanding the infection of individual microbial cells and the related biochemical changes remains limited. Using Raman microspectroscopy in conjunction with univariate and unsupervised machine learning approaches, we established a marker for identification of infected Gram-positive and Gram-negative bacteria. This non-destructive, label-free analytical method at single-cell resolution paves the way for future studies geared towards analyzing complex biological changes of virus-infected cells in pure culture and natural ecosystems.

## Introduction

Prokaryotic viruses substantially influence global ecosystems and biogeochemical cycles by infecting host populations. This predation can cause release of organic carbon and also enhance horizontal gene transfer (1), as viruses can act as mobile genetic elements (MGEs). Viruses are generally differentiated based on the type of genetic information stored in their viral particle - either single or double-stranded DNA or RNA (2). Viruses are also categorized based on their reproduction cycle as lysogenic or lytic (although seldomly other strategies like chronic infection or pseudolysogeny have been reported (3)). Viruses can insert their genetic information into the genome of an infected host and proliferate along with host reproduction (lysogeny). A lytic strategy involves the reorganization of host metabolism envisaging reproduction of viral particles and ultimately cell lysis. A host cell infected with a lytic virus is referred as a virocell and needs to be differentiated from ribocells, cells that generally proliferate irrespective of an infection (4). In a recent study, transcriptomics and proteomics were used to investigate whether metabolic differences between uninfected cells and virocells can impact an entire ecosystem (5). However, the study of virocells necessitates techniques that can capture virocell characteristics at the single cell level prior to cell lysis.

The development of confocal Raman microspectroscopy has enabled the measurement of single microbial cells (6), which consequently opened the possibility to gain insights into the heterogeneity of microbial communities (7). The combination of Raman microspectroscopy instruments with multivariate data analysis of digitally recorded spectra allowed for further increases in sensitivity in the last two decades, resulting in the detection of biochemical differences between bacterial species across growth phases (8). In this context, multivariate statistical analysis of Raman spectra has been used to differentiate single cells based on discrete wavenumbers corresponding to biochemical compounds. Huang and colleagues described a correlation between the fraction of ^13^C in the carbon source and a ratio shift based on Raman peaks of unlabeled ^12^C phenylalanine and ^13^C labeled phenylalanine (9). The ratio between isotopically labeled and unlabeled molecules can be applied to identify key degraders in mixed cultures and allows specific cell sorting for single cell methods (10). However, this sensitivity is also the bottleneck of this technique, as demonstrated by Garcia-Timmerman, who highlighted the influence of the sample preparation on the recorded spectra (11). Such differences complicate the construction of a public database and comparability of spectra across studies. Nevertheless, the comparison of selected wavenumbers between individual spectra of a single study is crucial for expeditious categorization of single cells based on their chemical composition.

In this study, the high sensitivity of Raman microspectroscopy to identify and characterize microstructural intracellular changes as well as viruses and their effects on host metabolism was harnessed to test whether this technology can be used in differentiating regular cells from virocells. To this end, three different model host-virus systems including DNA and RNA virus were used to analyze and monitor chemical changes during infection processes at single-cell level using a Raman microspectroscope. Multivariate statistics and unsupervised machine learning suggested identical wavenumbers contribute to differences between virocells and regular bacterial cells across our model systems. These wavenumbers were attributed to the ratio of nucleic acids and proteins, respectively, and are thus in congruence with current literature (5, 12, 13) regarding the differences of expression profiles of virocells.

## Results

### Hundreds of spectra acquired for individual cells display significant differences in the chemical composition of infected and uninfected cultures of P. syringae

Addition of *phi6* to *P. syringae* cultures resulted in the expected decline in optical density, enabling us to harvest cell populations representing a mixture of virocells and uninfected cells (Fig. 1A). We used this cell population and a culture without phage addition for comparison in single cell Raman microspectroscopy. In doing so, we successfully measured 448 high-quality spectra of individual *P. syringae* cells, of which 198 cells were measured after addition of *phi6*. The other 250 spectra were reference spectra from uninfected cells of *P. syringae*. Inspection of the spectra and comparison to previously published Raman spectra of bacteria confirmed the typically expected peaks for biomolecules confirming the measurement of actual microbial cells (8).

**Figure 1.**
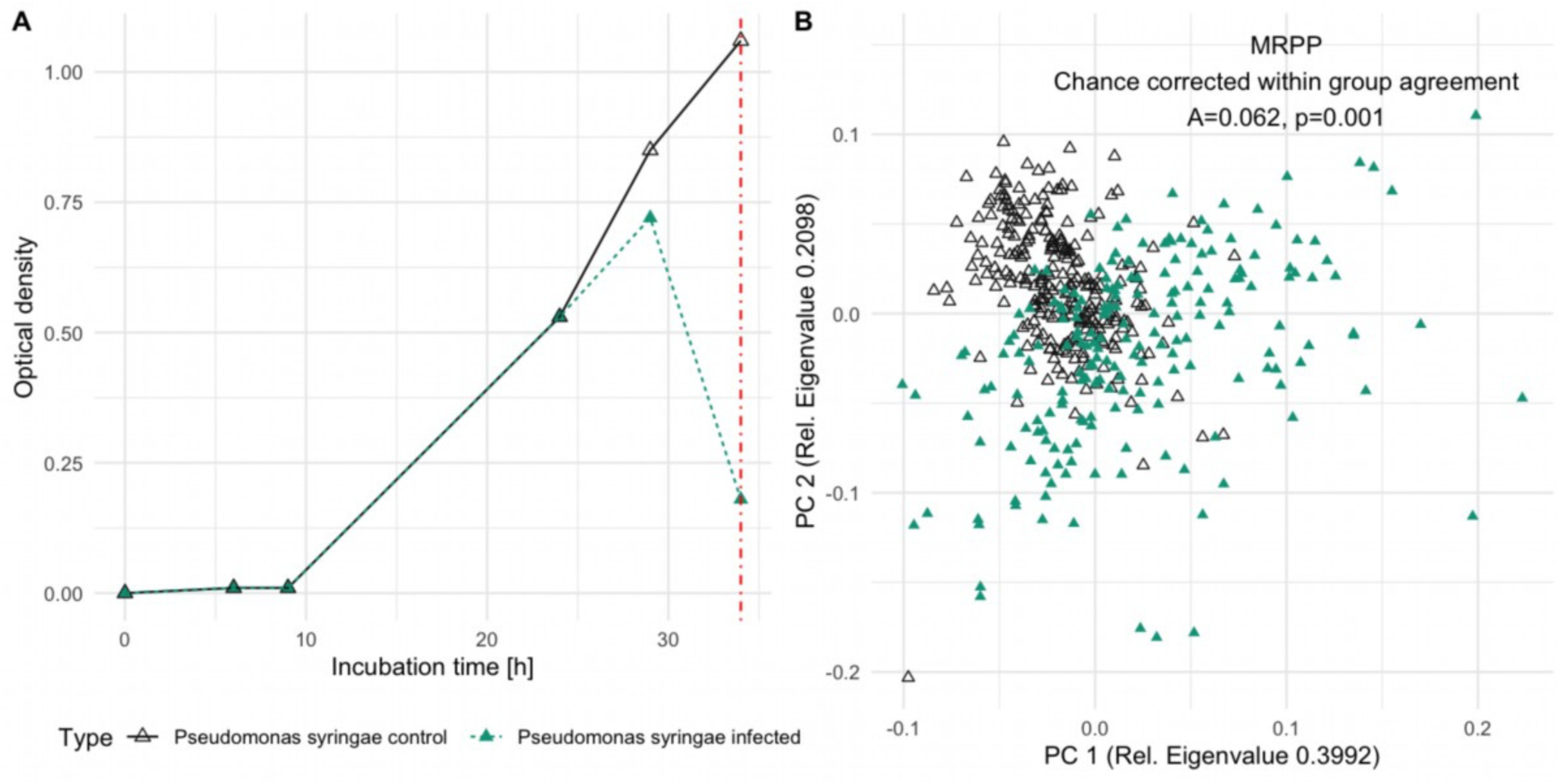
*Pseudomonas syringae* cultures without (black, empty triangles) and with (green, filled triangles) addition of phage phi6. The drop in optical density corresponds to viral cell lysis after 34 h; A: Growth curve determined by optical density; vertical red line highlighting the time of harvest. B: Principal Component Analyses of *P. syringae* single cell Raman spectra after lysis. (Ordination analyses based on Euclidean distance and spectral contrast angle revealed nearly identical results, see Figure S1) and the result of the Multi Response Permutation Procedure (MRPP) for control sample vs. infected sample.

Using the individual spectra of each measured cell, we computed an ordination analysis comparing individual cells of cultures with and without phage addition, which showed substantial differences (Fig. 1B). Importantly, the two datasets (with and without phage addition) were not entirely separated along Principal Component 1 (PC 1) or PC 2 but showed differences along both PCs, which agrees with the abovementioned mixture of virocells and uninfected cells in populations after phage addition. MRPP analysis displayed a highly significant p-value (<0.001), with a chance-corrected within group agreement (A) of 0.062. Consequently, phage addition and infection showed a significant and substantial change in the (bio)chemical composition of individual cells resulting in virocells.

To challenge the results of the observed differences between cultures with and without phage addition, we applied the abovementioned multivariate analysis to two different time points of the same uninfected culture of *P. syringae*. This experiment was set out with the aim of testing the null hypothesis that the differences between amended and non-amended cultures originate from variation during growth phases of cultures, which is known to exist in bacteria (8). The respective PCA (Fig. S2), which also includes the data from the amended cell culture, displays a difference of two timepoints along PC 2. However, the intragroup dissimilarity of the two individual timepoints was substantially lower than for the population amended with phage, particularly along the major component of the PCA. Although the MRPP testing for differences between the uninfected cultures at the two time points resulted in a significant p-value (0.002), the chance-corrected within group agreement was less than a sixth (0.009) of those identified for differences between cultures with and without phage addition. Moreover, comparing the combination of both time points of the uninfected culture to one with phage addition, we identified a highly significant difference (MRPP, p-value < 0.001, A = 0.06). Based on these observations the null hypothesis was rejected, supporting the working hypothesis that uninfected cultures can be distinguished from cultures with virocells using Raman spectroscopy.

### Differentiating wavenumbers of uninfected cells and virocells are attributable to nucleic acid and protein Raman shifts in P. syringae

To investigate the exact differences between cultures with and without phage addition as displayed in the PCA (Fig. 1B), we used the system of *P. syringae* / *phi6* for an in-depth statistical analysis. Comparing the contrast plot of phage-amended and non-amended cultures with the major two components of the PCA, highlighted the contribution of the individual wavenumbers that discriminate the two groups (cultures with and without phage addition; Fig 2A). Six wavenumbers contribute the most to the differences between the average spectra of the two groups with a high contribution to the PCA or a high density at the contrast plots and these are assigned to their respective biomolecules in Table 1.

**Table 1.**
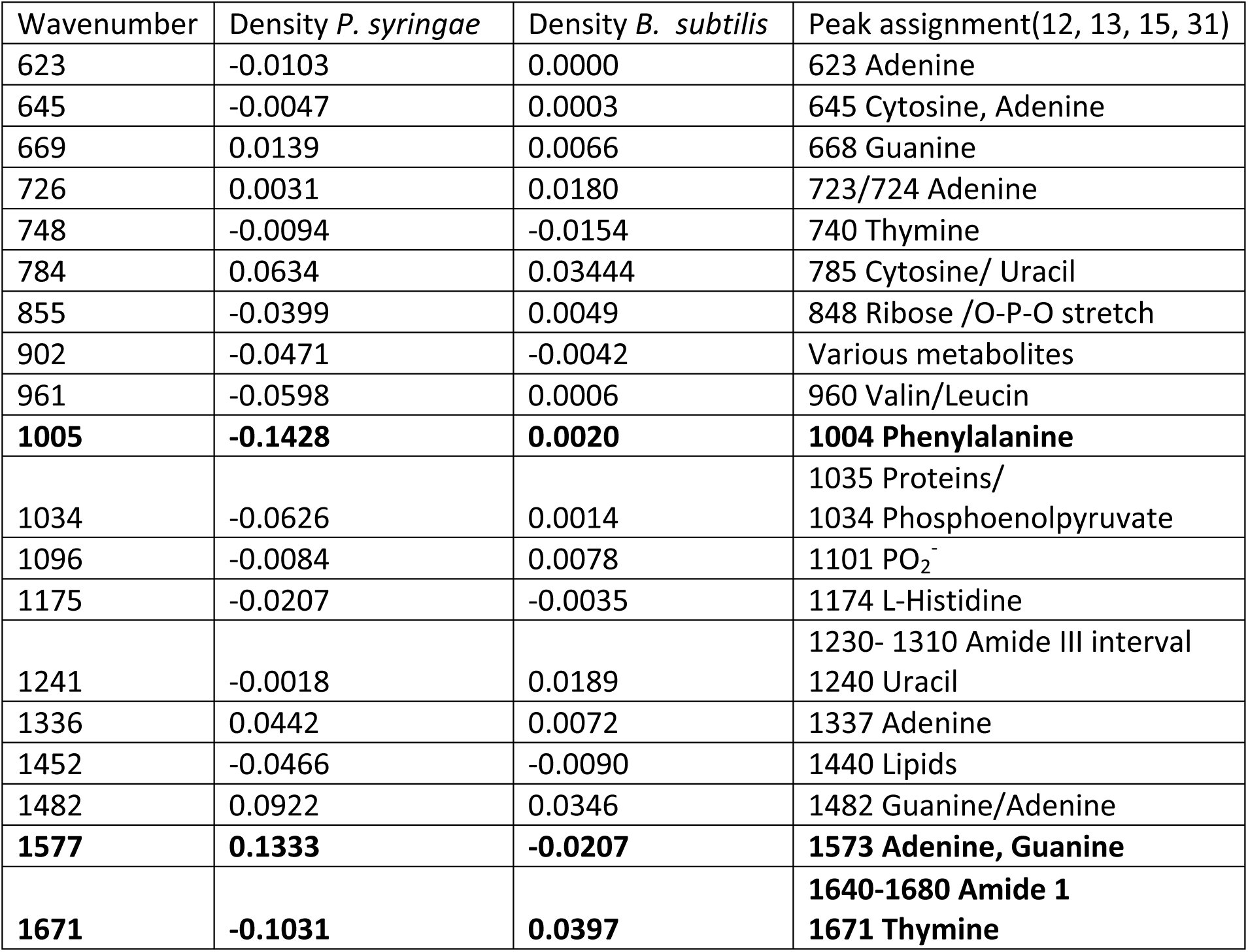
Wavenumbers assigned to biomolecules of microbial cells and their density in the contrast plots of infected and uninfected cells. A high positive density refers to prominence in infected cells, a negative value refers to wavenumbers more prominent in the control sample. Bold: Wavenumbers chosen for calculating the ratio for differentiating virocells from uninfected cells.

**Figure 2.**
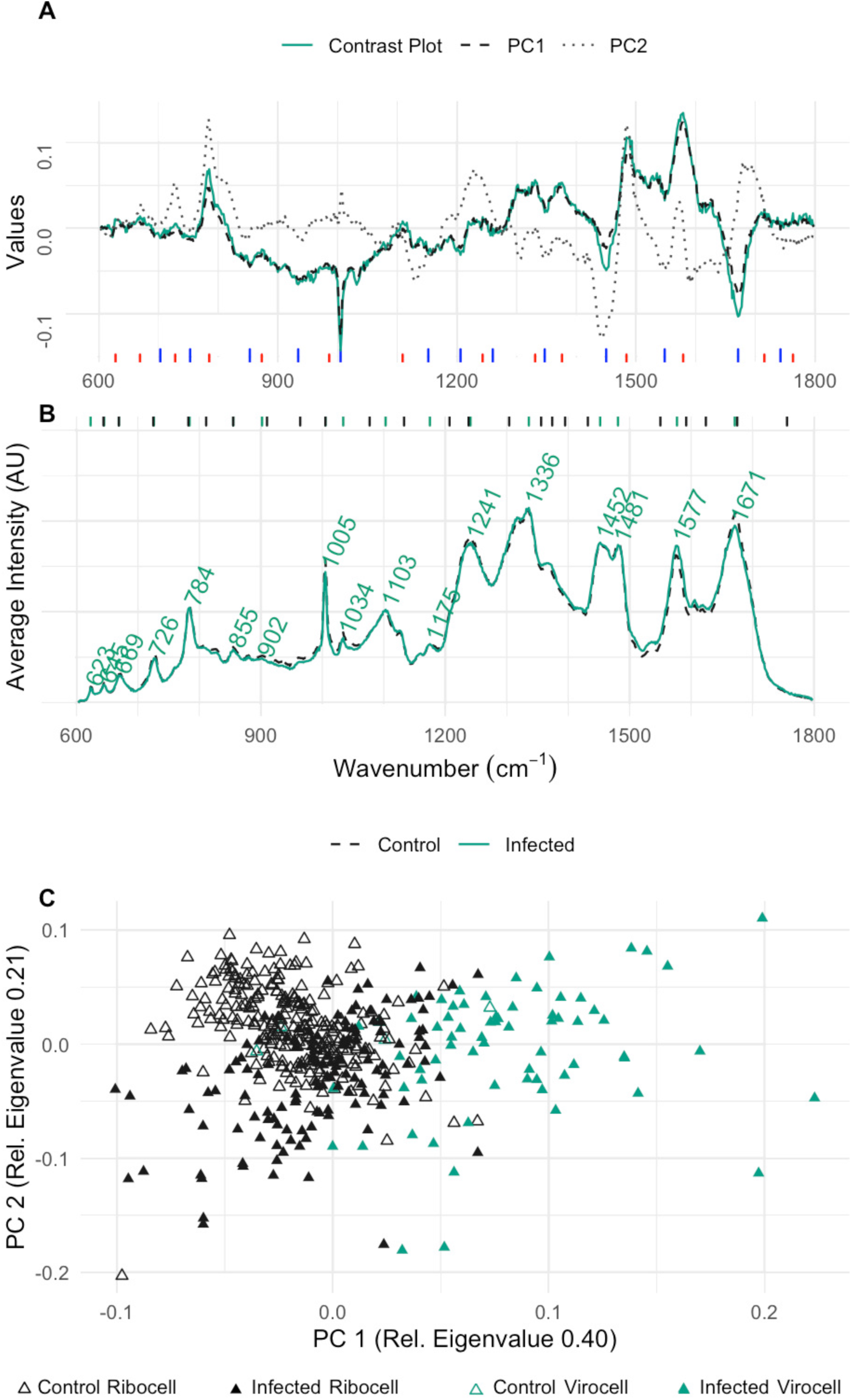
Evaluation of wavenumbers for virocell identification in *P. syringae*; **A**: Contrast plot (green) of potential infected cells in comparison to the wavenumber influence on PC 1 (black, dashed) and PC 2 (grey, line dotted), long, blue lines at the bottom indicate wavenumbers that decreased in virocells, red lines indicate wavenumbers increasing in virocells. **B**: Average Raman spectra of the samples with (green, solid) and without phage addition (black, dashed). Green lines at the top indicate the positions of the labeled peaks in the Raman spectra, black lines indicate peak maxima of the VIP of the OPLS; **C**: PCA of single cell Raman spectra from cultures with phages (filled triangle) and without phages (empty triangle); virocells identified based on the determined ratio are highlighted in green.

Three of the wavenumbers with the highest density in the contrast plot were assigned to nucleic acids (785, 1483 and 1576 1/cm), of which one was significantly higher in cultures amended with phage based on a Wilcoxon test (p_785_ = 0.15, p_1483_ = 0.71, p_1576_ = 2.2·10^−16^, respectively). By contrast, peaks assigned to proteins (1003 and 1671 1/cm) and lipids (1448 1/cm) are more prominent in the control sample, and the corresponding p-values of the proteins were significant (Wilcoxon test, p_1003_ = 1.7·10^−15^, p_1671_ = 2.2·10^−16^, p_1448_ = 0.71, respectively). Noteworthy, the wavenumbers associated with (highly) significant changes (1003, 1576 and 1671 1/cm) contributed to PC 1, while the other three with insignificant changes between phage amended and non-amended contributed more to PC 2 (785, 1440 and 1483 1/cm) (Fig. 2A). The intensities (I) of three wavenumbers with significant p-values were used to determine a differentiator for univariate differentiation of *P. syringae* virocells from uninfected cells (equation 1):

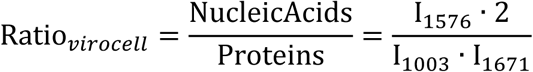

with wavenumbers assigned to proteins (1003 and 1671 1/cm) in the denominator and the nucleic acid peak (1576 1/cm) in the numerator. The Shapiro test demonstrated that the ratios based on spectra of the control sample based on equation 1 were normally distributed (p-value = 0.07) and those of the infected sample were not (p-value = 5.99·10^−9^), which was expected since the latter represent a mixture of virocells and uninfected cells. The calculated confidence intervals indicate, that *P. syringae* cells of the control group do not exceed a ratio of above 1.06 (99% probability), while this threshold was indeed exceeded (with a probability of 45 %, 66 of 198 cells) in the sample after phage addition. Consequently, equation 1 can be used to identify potential virocells in cultures of *P. syringae* (Fig 2C).

### Validation of selected wavenumbers for virocell identification of P. syringae via VIP of the OPLS model indicates high influence by peak shoulders

The average Raman spectra of the control sample and the infected sample show clear differences in the intensity of prominent biomolecule peaks chosen for the ratio determination (Fig 2B). However, plotting the peak maxima of the VIP of the OPLS together with the average Raman spectra revealed that the differences in virocells are sometimes not only represented by the maximum of the peak but also by its shoulders. This is a fine detail that is overseen by just visually inspecting the average spectra. The peaks can be assigned to their biomolecular origin (chemical bond) since their position does not change with a change in the molecular environment. However, the width of the peak is dependent on the composition of the molecule surrounding the polarized bond (14). Although the intensity change cannot be determined between two groups, this approach confirms the selected wavenumbers for the determined ratio for virocell identification in *P. syringae*.

### Applicability of virocell identification across three different species

Based on the differentiating ratio determined for virocells and uninfected cells of *P. syringae* (equation 1), we tested its applicability to other microbial species by repeating the analysis performed with *P. syringae* for *B. subtilis* and *M. mazei*. We calculated the ratios (equation 1) for cultures with and without virus addition, which showed a significant difference for *P. syringae* and *B. subtilis* (p-value < 0.0001, Fig. 3). By contrast, only a trend was revealed for the *M. mazei* / MetSV system (p-value < 0.0649; Fig. 3) without visible differences in PCA and contrast plot analysis (Fig. S3).

**Figure 3.**
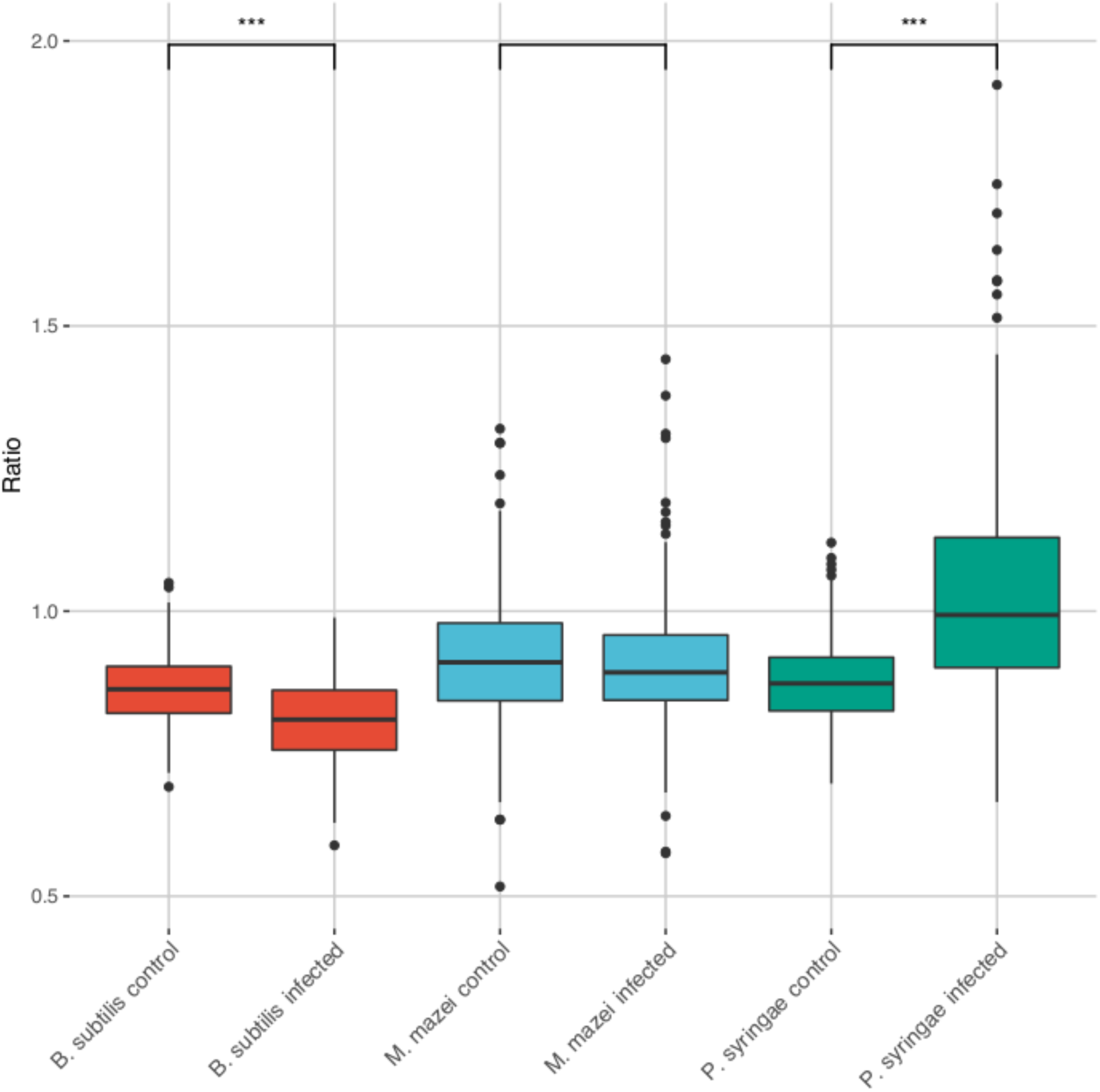
Boxplot of the determined ratio for control (no virus addition) and infected (with virus addition) samples of *B. subtilis* (red), *M. mazei* (blue) and *P. syringae* (green). Asterix indicating significance according to Wilcoxon Test (*** highly significant <0.0001, no asterix =not significant, *p-value < 0*.*06*). For a detailed multiple comparison across species (based on Dunn’s test) please see Supplementary Table 2.

For the *B. subtilis* / *phi29* system, a group of potential virocells could be differentiated from the control sample along PC 2 (Fig. 4C), yet the contrast plot (Fig. 4A) shows a lower density range than the one for *P. syringae* (Fig. 2A). The highest values contributing to spectra of infected cells were associated with nucleic acids and proteins (Wilcox test: p_785_ = 3.4·10^−11^, p_1483_= 3.0·10^−12^, p_1003_ = 0.15 and p_1671_ = 9.5·10^−12^), while peaks with the wavenumbers for hydrocarbons and nucleic acids were enriched in uninfected cells (p_1131_ = 1.7·10^−8^, p_1550_ < 2.2·10^−16^, p_1589_ < 2.2·10^−16^). Importantly, these identified wavenumbers included the same wavenumbers that were determined for the ratio (equation 1) for *P. syringae*. Although the associated signals of biomolecules were inverted compared to *P. syringae*, i.e., proteins were substantially higher in designated virocells and nucleic acids declined, the respective ratio (equation 1) can still be used to identify potential virocells of *B. subtilis* in Raman spectroscopy (Fig. 4).

**Figure 4.**
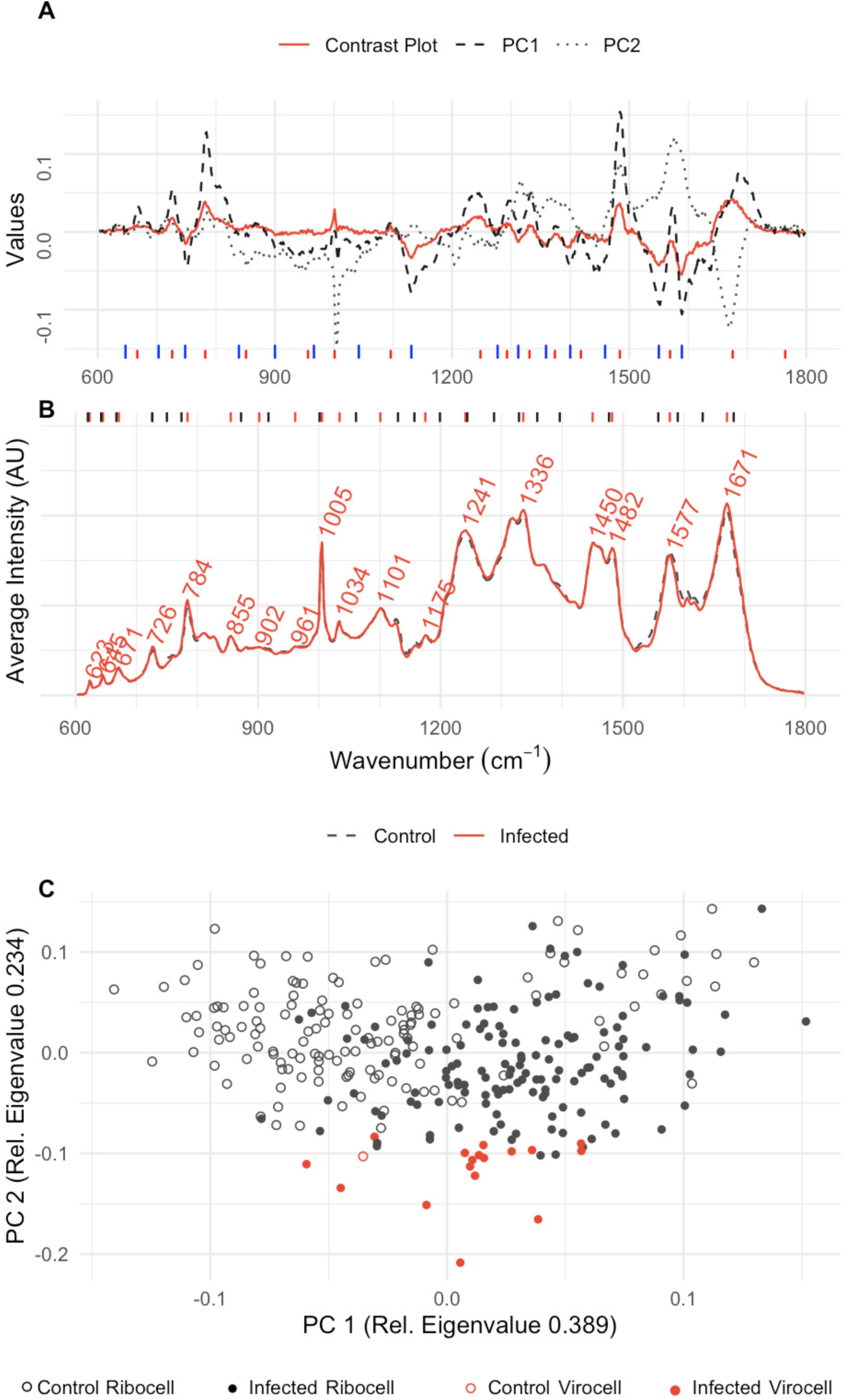
Evaluation of wavenumbers for virocell identification in *B. subtilis*; A: Contrast plot (red) of potential infected cells in comparison to the wavenumber influence on PC 1 (black, dashed) and PC 2 (grey, dotted), long, blue lines at the bottom indicate wavenumbers which decrease in virocells, red lines indicate wavenumbers increasing in virocells. **B**: Average Raman spectra of the samples with (red, solid) and without phage addition (black, dashed). Red lines at the top indicate the positions of the labeled peaks in the Raman spectra, black lines indicate peaks of the OPLS-importance; **C**: PCA of single cell Raman spectra with (filled dots) and without phage addition (empty dots), virocells identified based on the determined ratio are highlighted in red.

**Figure 5.**
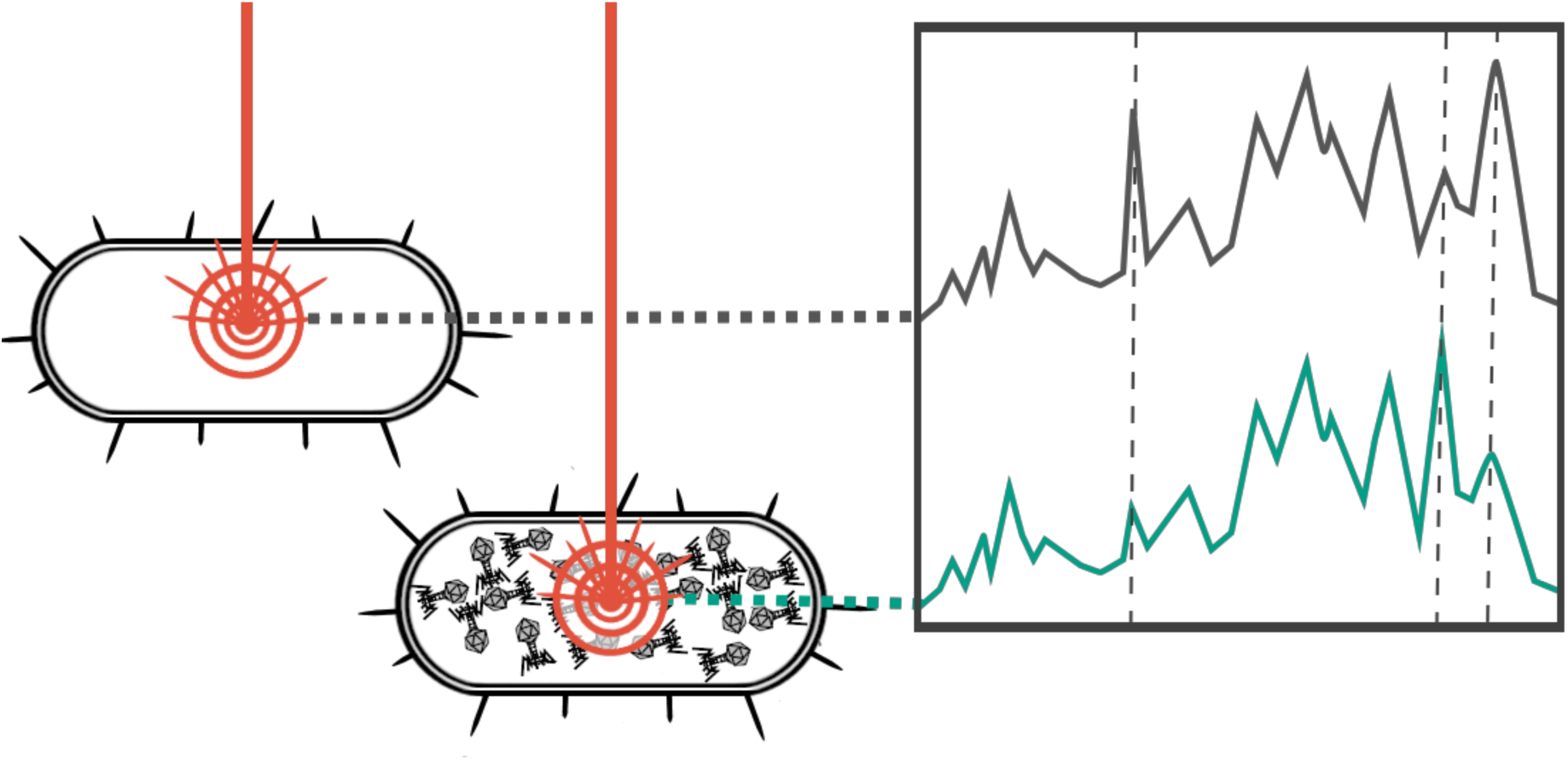
The laser of the Raman microspectroscope is focused on a single microbial cell. The presence of viral particles replicated inside the cell alters the Raman spectrum especially in the determined areas. Virocells can be determined by calculation a ratio based on these intensities.

## Discussion

In this manuscript we measured several hundred individual bacterial and archaeal cells (Fig. S4) to identify common changes in Raman spectra due to viral infections. One major challenge associated with measuring cultures of infected cells was their heterogeneity, meaning the culture consisting of uninfected cells and virocells at the same time. However, we were able to identify a specific ratio of Raman spectra that enabled us the differentiation of virocells and ribocells cells in the cultures of *P. syringae* and *B. subtilis*. This ratio was based on the wavenumbers 1003, 1576 and 1671 1/cm, which can be assigned to proteins and nucleic acid changes based on existing literature of recoded Raman spectra (12, 13).

### Overcoming challenges in identifying a Raman spectrum-based marker for virocells

For identification of a Raman spectrum-based marker of virocells, it was mandatory to use univariate and multivariate statistics (non-supervised machine learning) in concert. Neither univariate nor multivariate statistics alone were successful in identifying the respective wavenumbers necessary for the differentiation of virocells from uninfected cells. To initially identify a set of wavenumbers that showed differences between these two cell types we applied a multivariate analysis resulting in six wavenumbers, which were further filtered based on a Wilcoxon test to create the respective equation for differentiation of the two cell types. This was partly due to the fact, that multiple PCs can contribute to differences in statistical populations at various intensities, while we focused only on the two PCs with the greatest Eigenvalues. Two peaks of contributing substantially to PC 2 of both bacteria studied herein are assigned to guanine (1483 1/cm) (13) and the ring breathing of cytosine and uracil (785 1/cm) (12). Although this suggests a strong involvement of nucleic acid changes in uninfected vs. virocells, the Wilcoxon Test did not indicate a significant difference demonstrating an insufficient picture provided by multivariate data analysis (MRPP). On the other hand, using univariate statistics alone, the highest differences for the populations of *B. subtilis* did not occur at the maximum of the peak (1579 1/cm), which we determined from using both methods. Instead, the contrast plot had the highest values at the shoulders of the maximum peak, at 1550 1/cm and 1589 1/cm, suggesting that the peak position must be considered in Raman spectra via multivariate statistic. The reason for this phenomenon of the breadth of the peak can traced back to the chemical environment of the molecule as the Raman shift is characteristic for the polarized chemical bond. Several studies about differences of Raman spectra of packed and unpacked viral DNA/RNA and protein/oligonucleotide interactions have been performed in the past and describe altered base environments as reason for the observation of such perturbations (14–16).

### Phi29 likely causes a stress response in B. subtilis

The determined equation for differentiating virocells from ribocells in the *P. syringae*-*phi6* system, could also be applied to the *B. subtilis*-*phi29* system. However, the ratio used for the differentiation was significantly lower in the *B. subtilis* system, which is in stark contrast to the significantly higher ratio for *P. syringae*. The respective wavenumbers attributable to proteins (1003, 1671 1/cm) showed an increase in intensity in *B. subtilis*, and nucleic acids (1576 1/cm) appeared to decrease substantially during infection with *phi29*. A drop in nucleic acid content and increase in protein content (as observed here for virocells of *B. subtilis*) is complementary to multiple biological processes that can be observed for bacteria. Chemicals like ethanol can cause a similar change in the protein and nucleic acid content, which represents a stress response by the bacterium. This stress response was detected based on the same changes in the wavenumbers as observed here (17). However, the induction of lysogenic phage in *B. subtilis* was shown to result in a decrease of the Raman shifts at 782, 1095 1/cm and only a slight decrease at 1452, 1659 1/cm (18). The authors of the aforementioned study concluded that these measurements likely stem from the fact that the measured cell had ruptured, and an empty cellular hull had been measured (consisting of proteins and lipids, while nucleic acids are lost during lysis). They used Raman shifts around 1095 1/cm and 785 1/cm to measure the respective differences in the nucleic acid, both of which, did not show a significant difference in our datasets. Comparing these previous findings to our results for *B. subtilis*, someone can likely not differentiate between *B. subtilis* cells showing a stress response and a respective virocell. We conclude that *phi29* causes a stress response in *B. subtilis* during infection, which we measured during Raman spectra acquisition.

### High sensitivity of Raman spectra mirrors different types of phage infection

The changes in nucleic acid and protein content are contradictory in the *P. syringae* and the *B. subtilis* system and could not solely be attributed to complex stress responses, but rather to different types of phages. While *phi6* infecting *P. syringae* is a non-tailed RNA phage with a lipid membrane (19, 20), *phi29* is an DNA phage with a complex polypeptide structure consisting of a phage head and a phage tail (21). Consequently, an increase in the protein content during *phi29* replication can be associated with an increase in protein content in the cell. The wavenumber 1671 1/cm has previously been associated not only with the amides but also with thymine, a central component of DNA but not RNA (Table 1). Comparing the RNA phage *phi6* and the DNA phage *phi29*, we did observe a difference at the thymine concentration at this wavelength. A similar trend (increase in thymine/protein concentration) was also observed for the *M. mazei* system, which is also based on a DNA virus. We conclude that the putative increase of proteins measured at 1671 1/cm stems from an increase in protein and thymine concentration at the same time, reflecting the difference in DNA and RNA phage used in the experiments. Beyond the different types of phages, the relatively slow maturation of the *phi6* viral particles usually encompasses two different stages within the *P. syringae* host: After 45 min 50-nm particles can be observed within the host, and after 80 min these particles are covered by the viral membrane (22). The plot of the PCA in Fig 1 shows that infected cultures of *P. syringae* differed along both components. Component one was used for the ratio determination, but the ratio did not include spectra of individual virocells that showed a difference along component two. The shift of these virocells along PC2 was associated with a single wavenumber at 1448 1/cm. This wavenumber indicates an increase of lipids, which agrees with the production of lipid membranes for viral particle maturation (22). Consequently, we did not only succeed in identification of virocells of *P. syringae*, but also distinguishing the two infection stages during *phi6* maturation based on our Raman spectra.

## Conclusion

Our data encompassing 1,287 Raman spectra acquired for individual cells of three different microbial species with and without virus addition suggests that at least bacterial virocells can be differentiated from uninfected cells. We present a ratio of three wavenumbers that can be utilized to quickly perform this differentiation, although the type of phage (RNA vs. DNA) and different infection stages can influence the detection. Beyond detection, Raman spectra of individual cells are sensitive enough to capture essential information on the biology of individual phage-host systems. Namely, DNA and RNA phages, stress responses to the differentiation of maturation stages of phages within the microbial host cell can be robustly identified. We predict that the identification of such cells in batch culture experiments and ultimately in environmental samples will aid studying the biology of individual virocells and thus expand our understanding of the complex interplay of phage and hosts along with their associated biochemistry.

## Material and Methods

### Cultivation of model systems and sampling strategy

Two cultures of *Pseudomonas syringae* (DSM21482) were incubated at 25 °C with 150 rpm in Tryptone soya broth (DSM medium 545). After 24 h, the cultures reached the exponential growth phase and 1 vol% glycerol stock of the phage *phi6* (DSM21518) was added to one culture, the second culture was kept uninfected as negative control. Samples for Raman microspectroscopy were taken prior to phage addition and 10 hours after infection, indicated by a drop of the optical density.

*Bacillus subtilis* (DSM5547) was incubated at 37 °C with 150 rpm in DSM medium 545. After 4 h, the cultures reached the exponential growth phase and 10 vol% of a phage *phi29* solution (DSM5546) was added to one culture, the second culture was kept uninfected as control. The shaking was reduced to 80 rpm. Samples for Raman microspectroscopy were taken when the optical density dropped 2 h after infection (Fig. S5).

*Methanosarcina mazei* (DSM3647) was incubated and infected with Methanosarcina spherical virus (MetSV) as described previously (23). Samples for Raman spectroscopy were taken anaerobically before virus infection and 180 and 210 min after infection.

### Sample preparation for Raman microspectroscopy

Samples for Raman microspectroscopy were taken at respective time points from the model systems (see above). 1 mL of the culture was washed with 1 mL 1X PBS (pH 7.4, Sigma-Aldrich), followed by resuspension in 0.45 mL 1X PBS and 0.15 mL 4 % formaldehyde (Thermo scientific) solution (fixation at 4 °C for 3 h). Afterwards the sample was again washed in 0.5 mL 1X PBS and dehydrated at room temperature in 50 vol% and 80 vol % ethanol (Fisher Scientific) for 10 min each. Finally, the preparation was stored in 0.15 mL 96 % ethanol at -20 °C until spectral acquisition. Throughout all steps mentioned above, washing was done by pelleting of samples via centrifugation at 2,000 x g for 10 min, followed by discarding the supernatant.

### Raman spectral acquisition

Raman spectral acquisition was performed using a Renishaw in via™ Raman microspectroscope with a 532 nm Nd:YAG laser and 1800 l/mm grating, equipped with a Leica DM2700M microscope. A 100x dry objective with a numerical aperture of 0.85 was used. Daily calibration was performed using a silicon waver (Renishaw). For each dehydrated sample (preparation see above), a drop was placed on a highly polished steel slide (Renishaw) and air dried. For *Pseudomona*s *syringae* a spectral acquisition of 25-30 s at 10 % laser power and for *Bacillus subtilis* three accumulations of 25 s and 5 % laser power was used. For cells of *Methanosarcina mazei* a 15 s bleaching step prior to 30 s measurement at 5 % laser power was necessary to reduce the florescent background. At least 50 cells per drop were measured and at minimum three drops per sample were used.

### Multivariate statistical analyses

The spectra were imported to R (24) as SPC files and processed using R package *MicroRaman* (11). The spectral data were trimmed to a range of 600-1800 1/cm. After background subtraction using the Statistics-sensitive Non-linear Iterative Peak-clipping (SNIP) algorithm (25), data were normalized using Total Ion Current (TIC) (26). These preprocessed data were used to calculate Principal Components Analyses (PCA) (27)and dendrograms based on Euclidian distance (Ward D2 clustering) (24). PCA results were compared to Principal Coordinate Analyses (PCoA) (28) based on spectral contrast angle dissimilarities (11). Spectra of cells burnt during spectral acquisition, spectra of low intensity and those containing cosmic rays were identified and removed from the dataset. Wavenumbers causing differences between infected and uninfected spectra were identified using a contrast plot (11) and the influence on the principal components. Differences between the samples assessed via a Multi Response Permutation Procedure (MRPP) using 999 Monte Carlo permutations.

An orthogonal partial least square analysis (OPLS) (29) was performed on the baseline corrected data. The spectra were divided into “Species_control” or “Species_infected” according to the sample their originated from. The variable importance on projection (VIP) for each wavenumber in the range of 600 -1800 1/cm was determined and compared with the density of the contrast plot and the principal components.

The mean spectrum of each class was calculated by determining the mean intensity at each wavenumber.

### Determination of differentiating ratio of virocells and uninfected cells

Different combinations of the intensities of the three wavenumbers with the most influence in the contrast plots (contrasting virocells and uninfected cells) of the *P. syringae*-*phi6* system were further analyzed. The average intensities and the standard deviation were calculated for the normalized data of the uninfected cells and potential virocells. Then a Shapiro test for normal distribution was performed and a Wilcoxon test for non-normal distributed data was used to test if the data from infected and uninfected show a significant difference. For each ratio a cut-off value was defined to declare a cell as infected. The 99%-confidence interval was calculated for the infected group and the control group, afterwards the number of false positive spectra inside the control group was determined. The results derived from the *P. syringae*-*phi6* system for identification of differentiating wavenumbers was then applied to the other virus-host systems and a Dunn’s test was performed to differentiate between host type coupled to either infected/uninfected cultures (https://github.com/cran/dunn.test) (30).

In order to identify the respective Raman spectra and relate them to biomolecules, we followed various publications by G. J. Thomas and co-workers, which resulted in a collection of Raman spectra of nucleic acids and proteins (12) and De Gelder (13), who conducted a study on pure solutions of biomolecules. The assignments are summarized in Table 1.

## Acknowledgements

This work was supported by the DFG Grant PR1603/2-1. IM was supported by the Studienstiftung des deutschen Volkes, AJP acknowledges funding by the Ministry of Culture and Science of North Rhine-Westphalia (Nachwuchsgruppe “Dr. Alexander Probst”). We thank Sabrina Eisfeld and Agathe Materla for lab management and technical assistance. Priyanka Mishra and Rainer Meckenstock are acknowledged for support with the Raman microspectroscope.

